# A comparison of DNA/RNA extraction protocols for high-throughput sequencing of microbial communities

**DOI:** 10.1101/2020.11.13.370387

**Authors:** Justin P. Shaffer, Clarisse Marotz, Pedro Belda-Ferre, Cameron Martino, Stephen Wandro, Mehrbod Estaki, Rodolfo A. Salido, Carolina S. Carpenter, Livia S. Zaramela, Jeremiah J. Minich, MacKenzie Bryant, Karenina Sanders, Serena Fraraccio, Gail Ackermann, Gregory Humphrey, Austin D. Swafford, Sandrine Miller-Montgomery, Rob Knight

**Affiliations:** Department of Pediatrics, University of California, San Diego, La Jolla, CA; Center for Microbiome Innovation, University of California, San Diego, La Jolla, CA; Bioinformatics and Systems Biology Program, University of California San Diego, La Jolla, CA; Marine Biology Research Division, University of California, San Diego, La Jolla, CA; Department of Computer Science and Engineering, University of California, San Diego, La Jolla, CA; Micronoma Inc. San Diego, CA

**Keywords:** DNA extraction, RNA extraction, microbiome, microbial community, high-throughput sequencing, 16S rRNA, shotgun metagenomics, limit of detection, well-to-well contamination

## Abstract

One goal among microbial ecology researchers is to capture the maximum amount of information from all organisms in a sample. The recent COVID-19 pandemic, caused by the RNA virus SARS-CoV-2, has highlighted a gap in traditional DNA-based protocols, including the high-throughput methods we previously established as field standards. To enable simultaneous SARS-CoV-2 and microbial community profiling, we compare the relative performance of two total nucleic acid extraction protocols and our previously benchmarked protocol. We included a diverse panel of environmental and host-associated sample types, including body sites commonly swabbed for COVID-19 testing. Here we present results comparing the cost, processing time, DNA and RNA yield, microbial community composition, limit of detection, and well-to-well contamination, between these protocols.

**Accession numbers:** Raw sequence data were deposited at the European Nucleotide Archive (accession#: ERP124610) and raw and processed data are available at Qiita (Study ID: 12201). All processing and analysis code is available on GitHub (github.com/justinshaffer/Extraction_test_MagMAX).

**Methods summary:** To allow for downstream applications involving RNA-based organisms such as SARS-CoV-2, we compared the two extraction protocols designed to extract DNA and RNA against our previously established protocol for extracting only DNA for microbial community analyses. Across 10 diverse sample types, one of the two protocols was equivalent or better than our established DNA-based protocol. Our conclusion is based on per-sample comparisons of DNA and RNA yield, the number of quality sequences generated, microbial community alpha- and beta-diversity and taxonomic composition, the limit of detection, and extent of well-to-well contamination.

## Introduction

Our growing understanding of microbial communities continues to reveal knowledge important for fostering human and environmental sustainability [1–4]. Nearly every day new links are made between the human microbiome and human health [5–7], and the development of methods related to studying microbial communities is ever-expanding [8–10]. One major roadblock to studying microbial communities is that single methods rarely capture information from all organisms in a sample, or from across diverse sample types [11–13].

The ongoing COVID-19 pandemic driven by SARS-CoV-2 has infected over 40 million human individuals and killed 1.1 million (*i.e.*, as of October 18, 2020) [14]. Such an event represents an invaluable opportunity to study the effects of a novel pathogen on microbial interactions relevant to human hosts and other ecosystems [15–17]. Currently our protocols benchmarked for high-throughput microbiome sequencing focus on extracting high-quality DNA from samples [18], and therefore will not capture RNA-based genomes such as that of SARS-CoV-2, which is a positive-sense, single-stranded RNA virus [19].

Here, our aim was to identify an extraction protocol that extracts high-quality RNA, while also producing DNA output and community composition comparable to our previously benchmarked protocol [18]. We also considered technical differences regarding the detection ability [20] and extent of contamination [21–23], among protocols. We compared DNA and RNA yield, the number of quality sequences, microbial community alpha- and beta-diversity and taxonomic composition, the limit of detection, and extent of well-to-well contamination, across common sample types and among three extraction protocols.

## Materials & Methods

### Sample collection

To compare extraction protocols, we collected biological materials from a broad range of human- and environmental samples, focusing on types widely used in studies of microbial communities and SARS-CoV-2 detection [18,24,25]. Each unique sample was aliquoted across extraction plates for comparison of extraction efficiency among protocols. We included a total of 33 human skin samples, 30 human oral samples, 8 built environment samples, 6 fecal samples, 6 human urine samples, 6 soil samples, 4 water samples, 4 fermented food samples, and 2 tissue samples. We collected most sample types using Puritan wooden handled, cotton swabs, following the Earth Microbiome Project standard protocol [26]. To make comparisons relevant to SARS-CoV-2 detection, we collected additional samples mimicking those collected from patients using plastic handled, polyester swabs (BBL CultureSwab, Cat#: 220135; BD, Franklin Lakes, NJ), following the CDC’s specimen collection guidelines [24,25]

We collected samples such to allow for technical replication across three extraction protocols. Human skin samples included those from the foot, armpit, forehead, and nostril interior. Foot and armpit samples were collected from three individuals by rubbing five cotton swabs simultaneously on the sole of each foot or armpit, respectively, for 30 s. Forehead and nostril samples were collected from 12 individuals by rubbing two polyester swabs over the forehead for 30 s, or in each nostril for 15 s each, respectively. Human oral samples included throat, saliva, oral saline rinses, and the same rinses diluted in viral transport medium [27]. Throat samples were collected from 12 individuals by rubbing two polyester swabs across the pharynx for 30 s. Saliva was collected from 12 individuals by active spitting into a 50 mL centrifuge tube. Saline rinses were collected from three individuals by swishing 10 mL 0.9% saline for 30 s and spitting into a 50 mL centrifuge tube. To mimic storage in VTM, 5 mL of saline rinse was mixed with 100 μL 50X VTM in a 15 mL centrifuge tube. Built environment samples included floors and door handles. Floor and door handle samples were collected from two rooms using cotton swabs and two rooms using polyester swabs, by rubbing nine swabs simultaneously across the surface of a 1-ft^2^ tile for 30 s, or one entire door handle, respectively. Fecal samples included human, mouse, and cat samples. Human feces were collected from two individuals using commode collectors (Fisherbrand Commode Specimen Collection System). Mouse feces were collected from two individuals and stored in 1.5-mL microcentrifuge tubes. Cat feces were collected from two individuals and stored in plastic zip-top bags. Human urine samples included samples from female and male individuals. Urine was collected from three female- and three male individuals, and was stored in commode collectors or 50-mL centrifuge tubes. Soil samples included both tree rhizosphere- and bare soil. For each type, soil was collected from two adjacent sites down to a depth of 20 cm using a sterile trowel, and stored in plastic zip-top bags. Water samples included both fresh- and seawater, collected from two sites at the San Diego River, and two sites at Scripps Institute of Oceanography, respectively. Water was collected and stored in 50-mL centrifuge tubes. Fermented food samples included yogurt and sauerkraut samples. Two varieties of one brand of each food were purchased commercially and stored in 50-mL centrifuge tubes. Tissue samples included jejunum tissue from eight mice individuals: *ca.* 3.8 cm of the middle small intestine was removed, and any fecal matter inside was squeezed out lengthwise; each tissue section was added to a 2mL microcentrifuge tube containing 1mL sterile 1X PBS and *ca.* 40mg sterile 1-mm silicone beads, and homogenized at 6,000 rpm for 1 min with a MagNA Lyser (Roche Diagnostics, Santa Clara, CA). The liquid homogenate from three intestinal sections from cohoused mice was pooled to create a single sample (one sample per cage). All samples were stored at −80°C within 3 h of collection, and were frozen for a maximum of 24 h before extraction. To compare limits of detection, we included serial dilutions of cultures of *Bacillus subtilis* (Firmicutes) and *Paracoccus denitrificans* (Alphaproteobacteria) [20]. Input cell densities ranged from 2.0-9.6E7 cells for *B. subtilis,* and 0.0-3.1E7 cells for *P. denitrificans*. To compare well-to-well contamination (*e.g.*, Minich et al. 2019), we included plasmid-borne, synthetic 16S rRNA gene spike-ins (*i.e.*, 4ng of unique spike-in to one well of each column in the plate) [28], and at least five extraction blanks per plate.

### DNA and RNA extraction

We compared two extraction protocols that use a 96-sample, magnetic bead cleanup format: the Qiagen MagAttract PowerSoil DNA Isolation Kit (Cat#: 27000-4-KF; Qiagen, Carlsbad, CA), and the Applied Biosystems MagMAX Microbiome Ultra Nucleic Acid Isolation Kit (Cat#: A42357; Applied Biosystems, Foster City, CA). We considered that the PowerSoil kit protocol includes heating the lysis solution to 60°C when mixing with samples, and a subsequent 20-min bead beating step, whereas the MagMAX kit has no heating and only a 2-min bead beating step. Additional heating and extended bead-beating may alter the extent of cellular lysis and degradation of extracellular nucleic acids, and subsequently microbial community composition. We therefore included a third protocol: a variant of the MagMAX one including 60°C incubation and 20-min bead beating steps, and refer to the three protocols as “PowerSoil”, “MagMAX (20-min)” and “MagMAX (2-min)”.

For extraction, aliquots of each sample were transferred to unique wells of a 96-well extraction plate. For samples collected with swabs, the entire swab head was broken off into the lysis plate. For liquid samples, we transferred 200 μL. For bulk samples, we used cotton swabs to collect *ca.* 100 mg of homogenized material and broke the entire swab head off into the lysis tube. Extractions were performed following the manufacturer’s protocol, except for the modifications made to the MagMAX (20-min) protocol described above. Lysis was performed with a TissueLyser II (Qiagen, Carlsbad, CA). Bead clean-ups were performed with the KingFisher Flex Purification System (ThermoFisher Scientific, Waltham, MA). Extracted nucleic acids were stored at −80°C prior to quantification of RNA yield, fragment length distribution, and integrity, as well as quantification of DNA yield and downstream sequencing.

### 16S rRNA gene and shotgun metagenomics sequencing

We prepared DNA for 16S rRNA gene- and shallow shotgun metagenomics sequencing as described previously [10,29–31]. For 16S data, raw sequence files were demultiplexed using Qiita [32], and sub-operational taxonomic units (sOTUs) generated using Deblur [33]. For shallow shotgun metagenomics data, raw sequence files were demultiplexed using BaseSpace (Illumina, La Jolla, CA), quality-filtered using Atropos [34] and human read depleted by alignment to human reference genome GRCh38 using bowtie2 [35]. Filtered reads were aligned to the Web of Life database [36] using Shogun [31] with default parameters and using bowtie2 as the aligner, followed by read classification with the Web of Life Toolkit App [36,37]. Raw sequence data were deposited at the European Nucleotide Archive (accession#: ERP124610) and raw and processed data are available at Qiita (Study ID: 12201). All processing and analysis code is available on GitHub (github.com/justinshaffer/Extraction_test_MagMAX).

## Results & Discussion

We found DNA yield to be similar across the three extraction protocols, and note that when considering all sample types (*n* = 615 samples), the extraction efficiency of the PowerSoil protocol was more similar to that of the MagMAX (20-min) one as compared to MagMAX (2-min) (paired-sample Wilcoxon signed-rank test: PowerSoil vs. MagMAX-20-min, *W* = 10540, *p* = 0.6); PowerSoil vs. MagMAX-2-min, *W* = 81170, *p* = 0.01) (Figure S1). We observed a similar pattern for the number of quality-filtered 16S reads (PowerSoil vs. MagMAX-20-min, *W* = 11482, *p* = 0.1; PowerSoil vs. MagMAX-2-min, *W* = 4651, *p* = 2.74E-11), however for quality- and human-filtered shotgun metagenomics reads, both MagMAX protocols varied from the PowerSoil one (PowerSoil vs. MagMAX-20-min, *W* = 15873, *p* = 1.41E-11; PowerSoil vs. MagMAX-2-min, *W* = 17148, *p* = 2.24E-15) (Figures 1 & S2).

**Figure 1.**
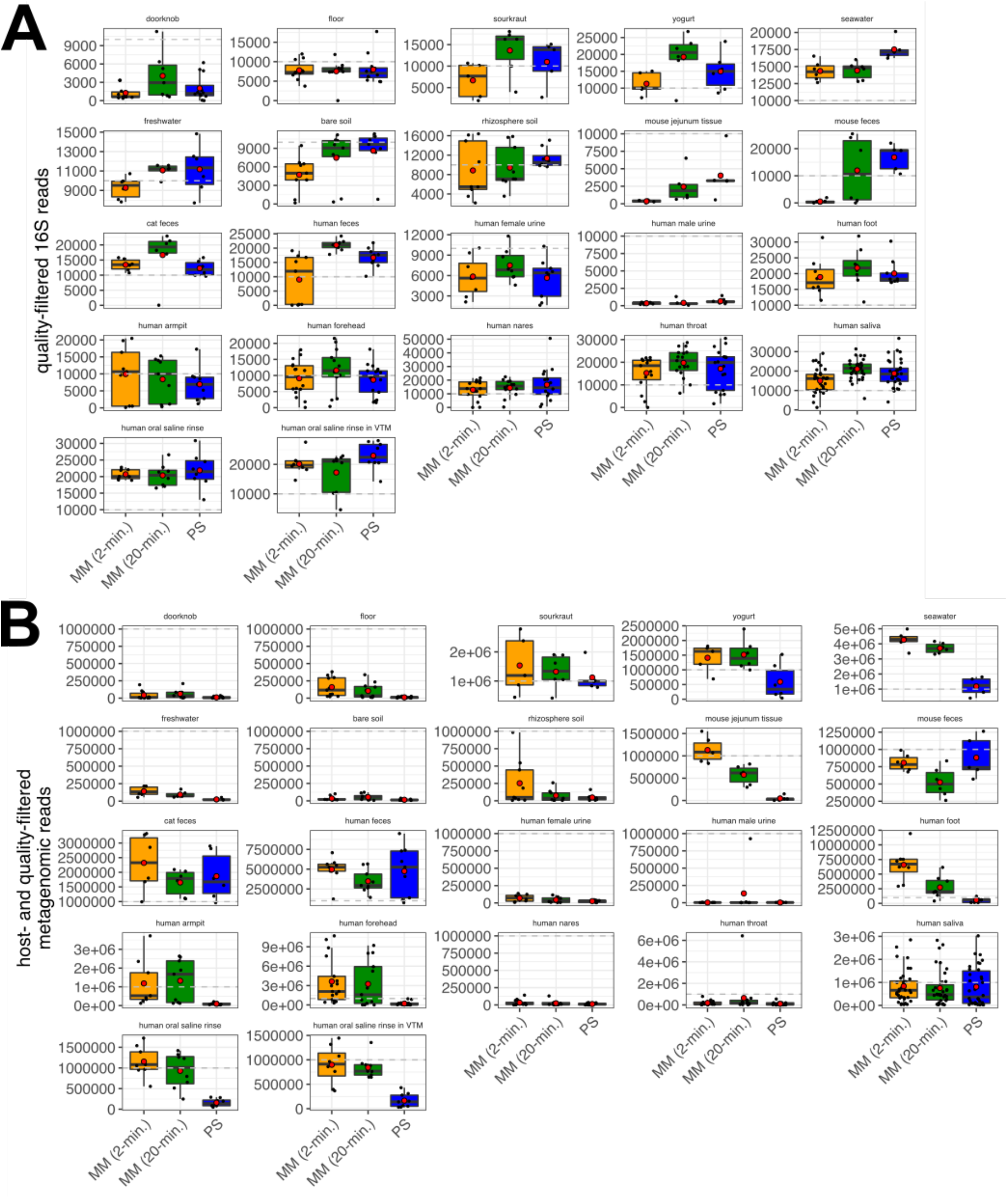
Average number of quality sequences for (1) 16S and (B) metagenomics data (*n* = 660 samples included). MM = MagMAX; PS = PowerSoil. Red circles indicate means. Dashed lines indicate our expectation of (A) 10,000 from 16S and (B) 1 million reads from metagenomics, respectively, for human fecal samples. Note that additional samples included here absent from our statistical test (*n* = 45) include those for which technical replication across protocols was not feasible due to recommended sampling protocols (e.g., human nares, human throat), so we included biological replicates instead. Sample types missing here lacked representation by both MagMAX protocols.

From a technical perspective, our comparison of the limit of detection (LOD) of each protocol indicates that the MagMAX (2-min) protocol requires 10X the number of cells required by PowerSoil for accurate detection in mixed bacterial cultures (Table 1). This is compared to 10,000X required by the MagMAX (20-min) protocol (Table 1). This pattern is mirrored when considering sample retention following filtering based on LOD thresholds, for which the MagMAX (2-min) is better with high-biomass samples and the MagMAX (20-min) with low-biomass ones. However, we observed an increase in well-to-well contamination in the MagMAX (20-min) protocol as compared to the MagMAX (2-min) one (Figure 2). This indicates that mimicking lysis parameters from the PowerSoil protocol in the MagMAX one has undesirable consequences, and that the MagMAX (2-min) protocol should be favored from this perspective.

**Table 1.** Limits of detection across extraction protocols. Titrations of cultured cells were used to identify the number of reads needed per sample, to meet various thresholds of detection (i.e., the percentage of reads mapped to expected taxa vs. background contaminants). Read depths corresponding to a threshold of 50% were used for filtering samples prior to community analyses of microbial 16S data, as recommended [20]. The retention of samples following filtering based on the read depth for each threshold is shown. Gram (+) = *Bacillus subtilis*; Gram (−) = *Paracoccus denitrificans*; mixed culture = *B. subtilis* and *P. denitrificans*.

| extraction protocol | sample set | threshold | LOD Gram (+) | LOD Gram (-) | LOD mixed culture | read depth | samples retained | % samples retained |
| --- | --- | --- | --- | --- | --- | --- | --- | --- |
| MagMAX (2-min) | High biomass | 50% | - | - | 5.73E+02 | 362 | 76 | 80% |
|  |  | 80% | - | - | 5.73E+04 | 1392 | 69 | 73% |
|  |  | 90% | - | - | 5.73E+04 | 3512 | 64 | 67% |
|  |  | 95% | - | - | 5.73E+06 | 9144 | 46 | 48% |
|  | Low biomass | 50% | 1.60E+03 | 3.10E+03 | - | 637 | 69 | 73% |
|  |  | 80% | 1.60E+04 | 3.10E+03 | - | 1631 | 63 | 66% |
|  |  | 90% | 1.60E+04 | 3.10E+03 | - | 3007 | 59 | 62% |
|  |  | 95% | 1.60E+05 | 3.10E+04 | - | 5526 | 52 | 55% |
| MagMAX (20-min) | High biomass | 50% | - | - | 5.73E+05 | 8499 | 68 | 71% |
|  |  | 80% | - | - | 5.73E+07 | 14522 | 55 | 57% |
|  |  | 90% | - | - | 5.73E+07 | 20158 | 30 | 31% |
|  |  | 95% | - | - | 5.73E+07 | 27541 | 3 | 3% |
|  | Low biomass | 50% | 1.60E+01 | 3.10E+01 | - | 491 | 79 | 83% |
|  |  | 80% | 1.60E+03 | 3.10E+03 | - | 776 | 71 | 75% |
|  |  | 90% | 1.60E+03 | 3.10E+03 | - | 1031 | 68 | 72% |
|  |  | 95% | 1.60E+04 | 3.10E+03 | - | 1354 | 64 | 67% |
| PowerSoil | High biomass | 50% | - | - | 5.70E+01 | 1050 | 87 | 92% |
|  |  | 80% | - | - | 5.73E+07 | 14632 | 36 | 38% |
|  |  | 90% | - | - | NA | 106110 | 0 | 0% |
|  |  | 95% | - | - | NA | 944308 | 0 | 0% |
|  | Low biomass | 50% | 1.60E+03 | 3.10E+00 | - | 1836 | 69 | 72% |
|  |  | 80% | 1.60E+04 | 3.10E+02 | - | 3797 | 62 | 65% |
|  |  | 90% | 1.60E+05 | 3.10E+03 | - | 5997 | 57 | 59% |
|  |  | 95% | 1.60E+05 | 3.10E+03 | - | 9345 | 40 | 42% |

**Table 2.**
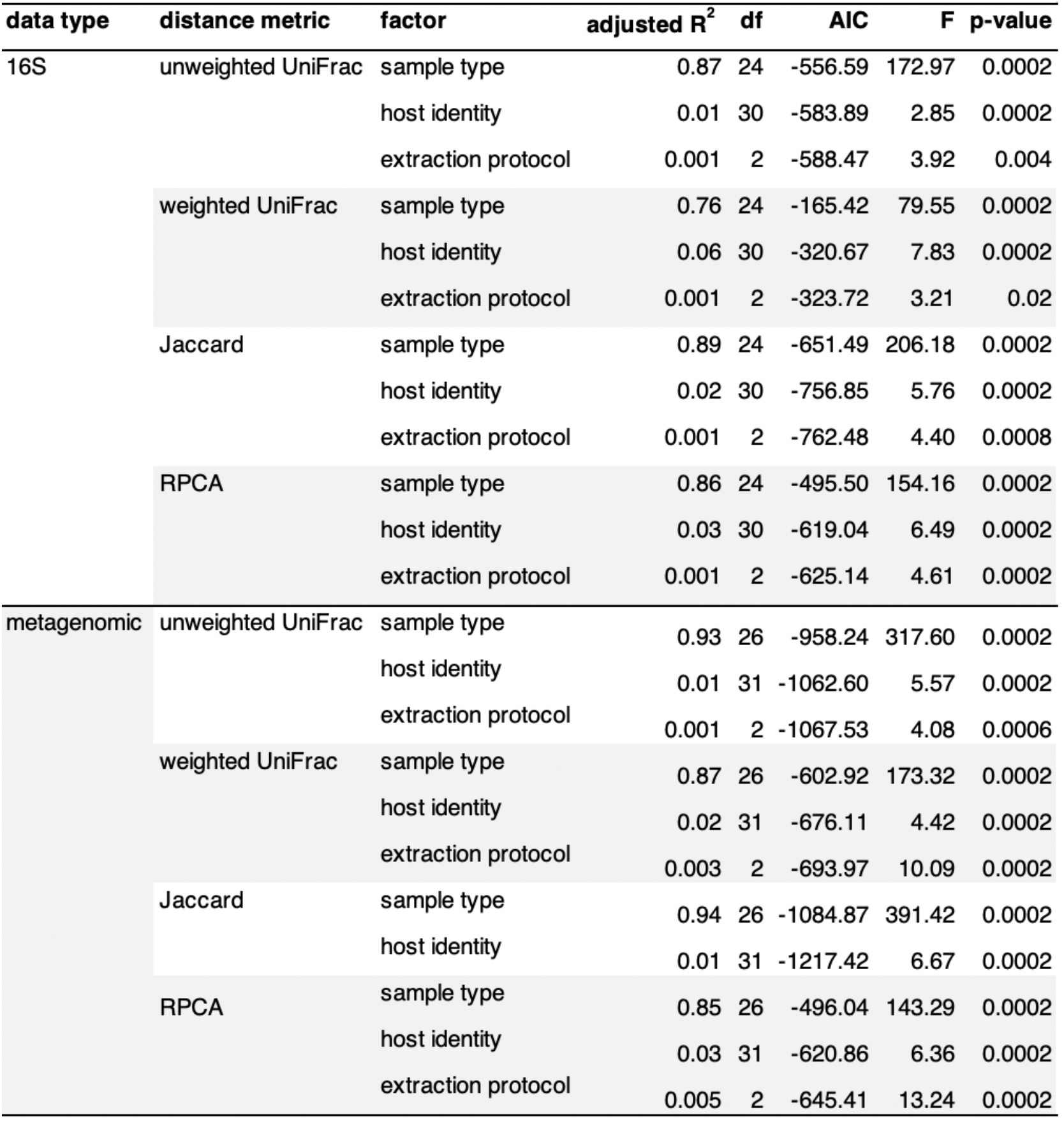
Results from a forward, stepwise model selection of factors influencing microbial community beta-diversity. Values are based on permutation tests of variation explained by redundancy analysis, done separately for four unique metrics for both 16S and metagenomics data. The full model included bead beating time (i.e., 2- vs. 20-min.), sample biomass (i.e., high- vs. low-biomass), sample type, host subject identity, and extraction protocol (i.e., MagMAX [2- min.], MagMAX [20-min.], PowerSoil), as model variables. 16S data were rarefied as noted for Figure 3. Metagenomics data were rarefied to 17,000 host- and quality-filtered reads per sample, or had samples with fewer than 17,000 reads excluded when using RPCA distances (*n* = 647 samples). Rarefaction depths were selected to maintain at least 75% samples from both high- and low-biomass datasets.

**Figure 2.**
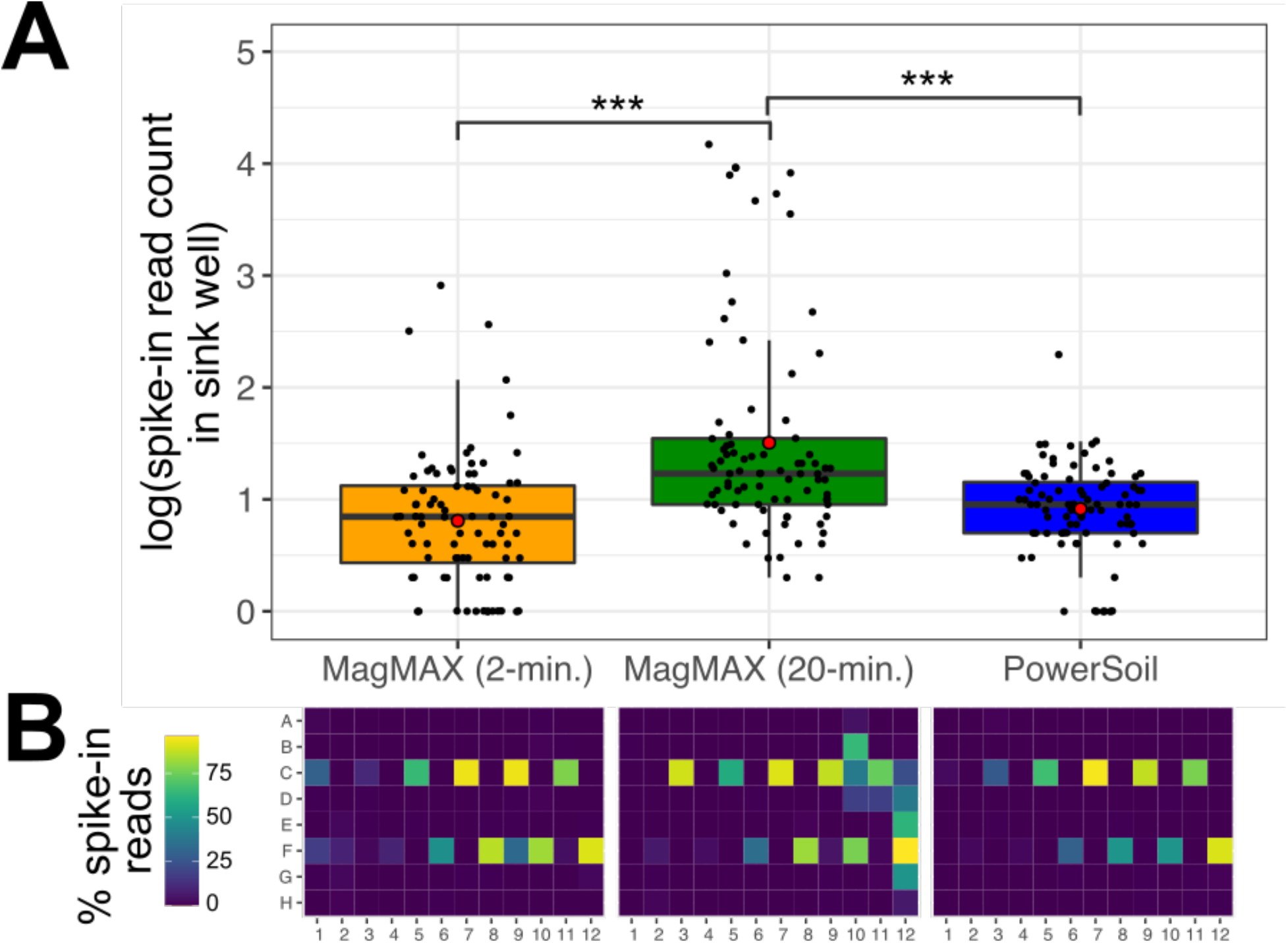
Well-to-well contamination among the three extraction protocols. Plasmids harboring synthetic 16S sequences were spiked into a single well per plate column (i.e., alternating from row C to F across columns: C1, F2, C3, F4, etc.) of each high-biomass sample plate prior to extraction. (A) The number of reads matching synthetic 16S sequences was quantified for all wells that did not receive a spike-in. Asterisks indicate significant differences between pairs of extraction protocols as determined by a Kruskal-Wallis post-hoc Dunn’s test with a Benjimini-Hochberg correction for multiple comparisons; *** *p* < 0.001. (B) The percentage of spike-in reads among all reads per well shown as a heatmap.

With respect to microbial community composition, we found bias introduced by extraction protocol to be small compared to variation among sample types or replicates of the same sample (*i.e.*, 1-2 orders of magnitude weaker in explaining beta-diversity; Table 1, Figures. S3 & S4). We also found strong correlations in microbial community beta-diversity among samples between any two extraction protocols, however relationships with the PowerSoil protocol were slightly stronger for MagMAX (2-min) as compared to the MagMAX (20-min) (Table S1). We used principal coordinates analysis (PCoA) of unweighted UniFrac distances to visualize these trends, and confirmed that samples clustered strongly by type and host subject and not by extraction protocol, for both 16S and metagenomics data (Figure 3; other distance metrics shown in Figures S5 & S6). Estimates of alpha-diversity were more comparable to those from PowerSoil for the MagMAX (2-min) protocol (paired-sample Wilcoxon signed-rank test: PowerSoil vs. MagMAX-2-min, *W* = 5916, *p* = 0.0001; PowerSoil vs. MagMAX-20-min, *W* = 7058, *p* = 1.53E-06) (Figures 4 & S7). Finally, the majority of genera (16S) and species (metagenomics) were shared across all three extraction protocols, however for both datasets the MagMAX (2-min) protocol shared a greater number of exclusive taxa with the PowerSoil protocol as than the MagMAX (20-min) protocol did (Figure 5).

**Figure 3.**
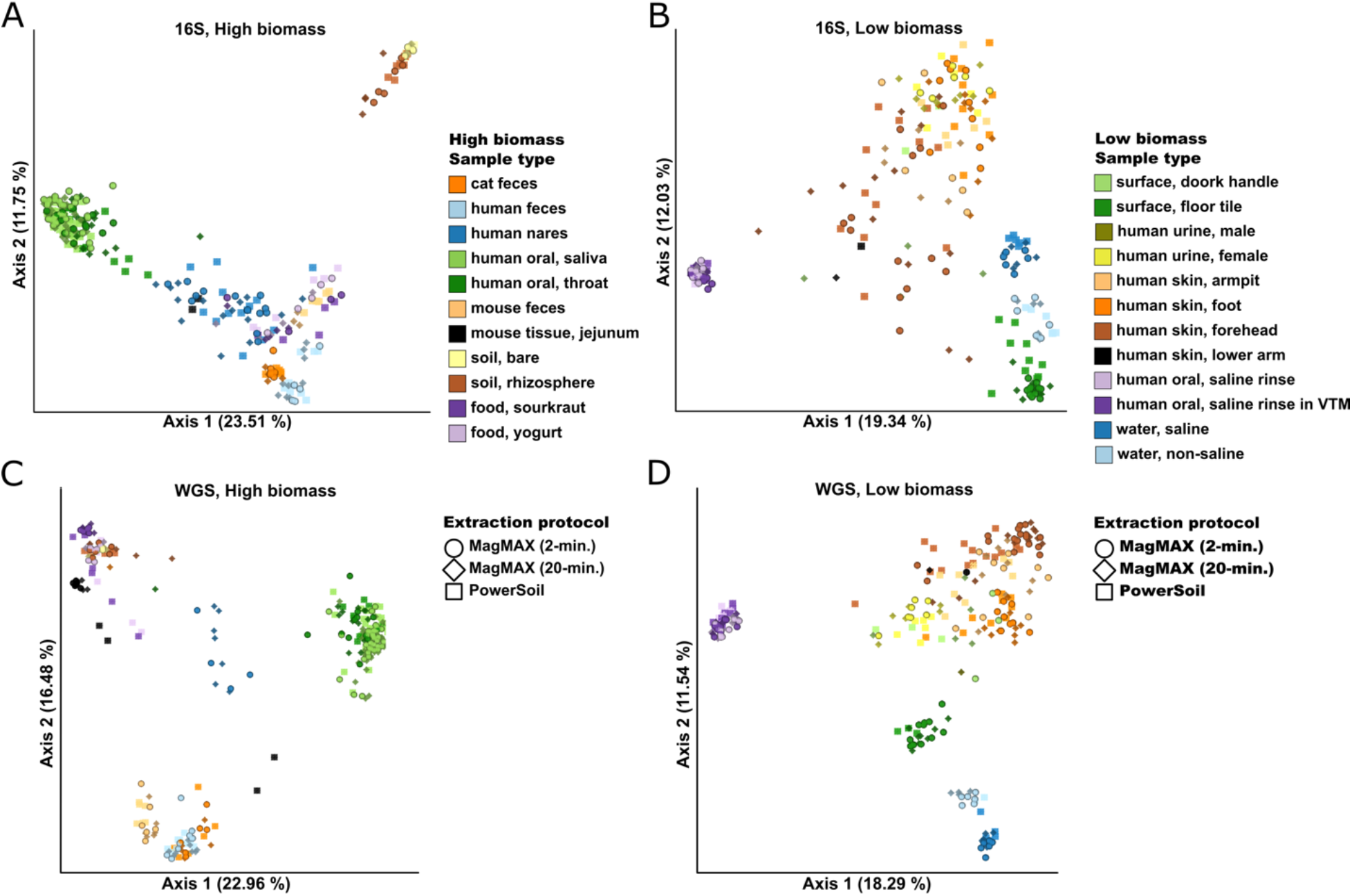
Principal coordinates analysis (PCoA) plots showing unweighted UniFrac distances based on 16S data for (A) high biomass samples and (B) low biomass samples, and shotgun metagenomics data for (C) high biomass samples and (D) low biomass samples. Colors indicate sample types and shapes indicate extraction protocols. Mock community and control blanks were excluded for clarity. 16S data were rarefied to 5,000 quality-filtered reads per sample for both high- and low-biomass samples (*n* = 611 samples). Metagenomics data were rarefied to 35,000 host- and quality-filtered reads per high-biomass sample (*n* = 287 samples), and to 20,000 reads per low-biomass sample (*n* = 242 samples). When using RPCA distances rather than using rarefied data, we excluded samples with fewer reads than the rarefaction depth for that dataset. Rarefaction depths were selected to maintain at least 75% samples from both high- and low-biomass datasets.

**Figure 4.**
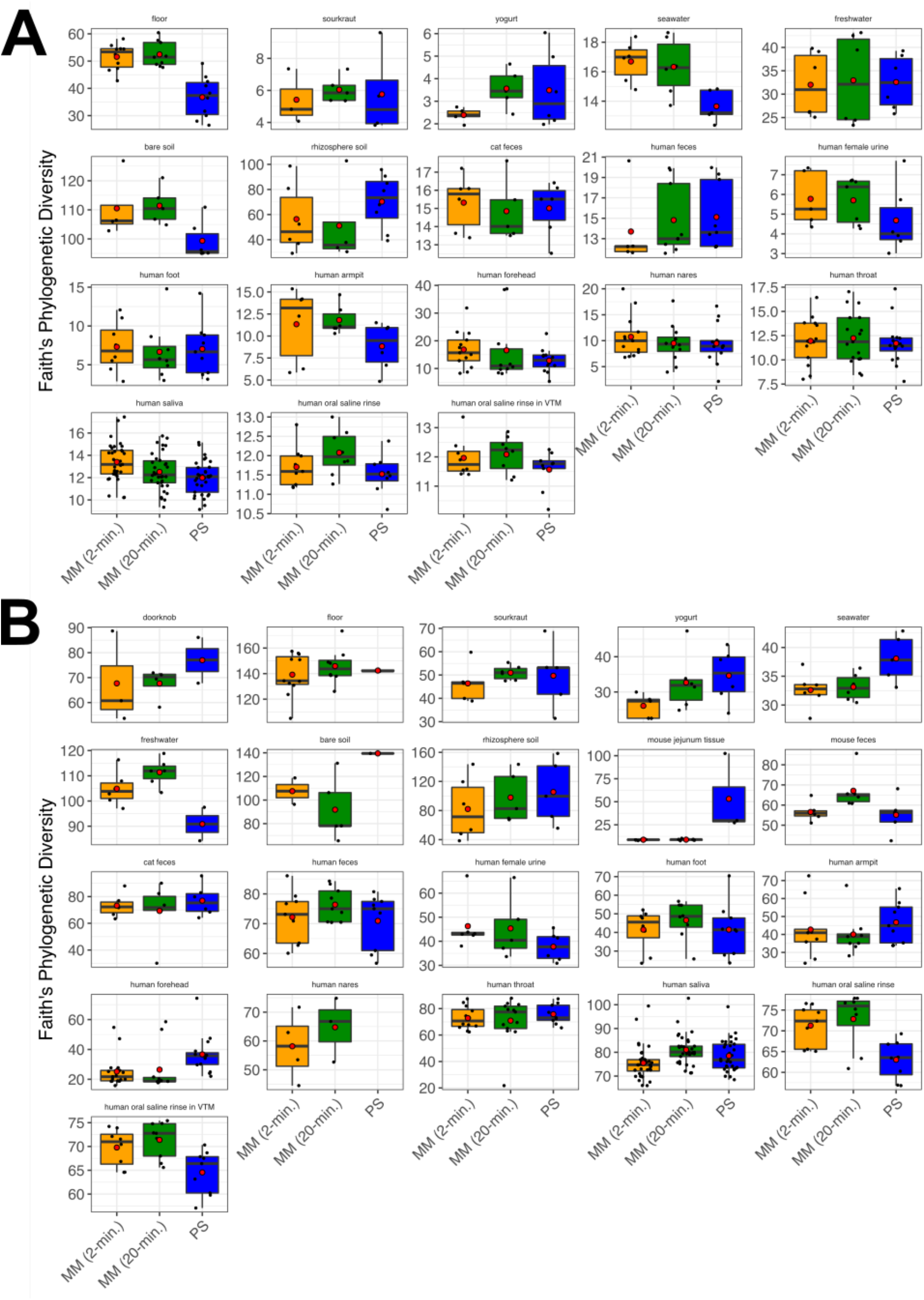
Alpha-diversity (Faith’s Phylogenetic Diversity) among the three extraction protocols based on (A) 16S and (B) metagenomics data. MM = MagMAX; PS = PowerSoil. Red circles indicate means. Data were rarefied as noted for Figure 3.

**Figure 5.**
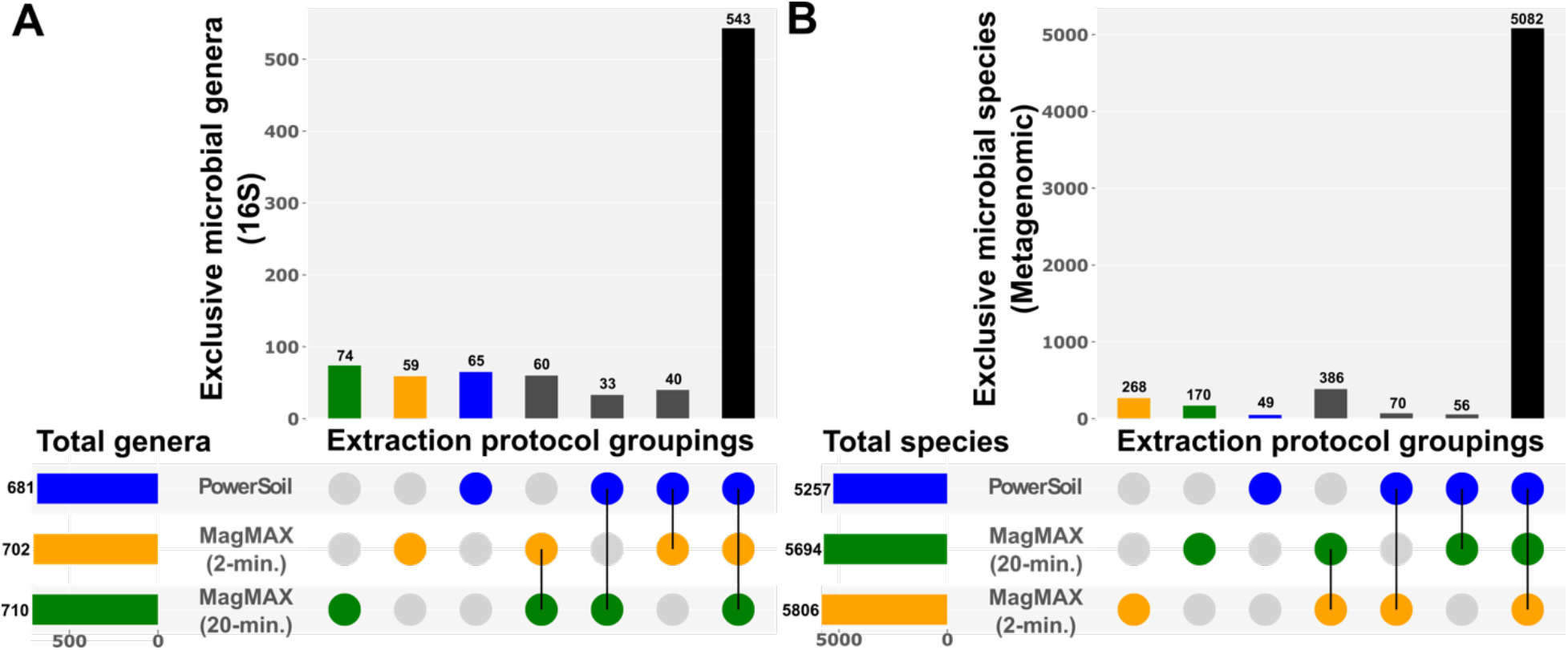
Upset plots showing (A) genera for 16S data and (B) species for metagenomics data, highlighting taxa shared among extraction protocols. Data were rarefied as noted for Figure 3.

Together, these results highlight that, despite variation in DNA yield, sequence read counts, and the limit of detection of microbial cells among extraction protocols, differences in microbial taxonomic- and community composition resulting from the different methods were minor, for both 16S and metagenomic microbial sequence data. However, between the two MagMAX protocols, we note that for beta-diversity, alpha-diversity, and taxonomic composition, the MagMAX (2-min) protocol generates more comparable results to the PowerSoil protocol.

Importantly, whereas RNA yield was comparable between the two MagMAX protocols (Figure 6A), we observed a higher quality of RNA extracted using the MagMAX (2-min) vs. the MagMAX (20-min) protocol (Figure 6B & C). In addition to reduced well-to-well contamination from a shorter bead-beating time during lysis for the MagMAX (2-min) vs. the MagMAX (20-min) protocol, the lack of incubation of the lysis buffer results in relatively high-quality RNA produced with the former vs. the latter (Figure 6).

**Figure 6.**
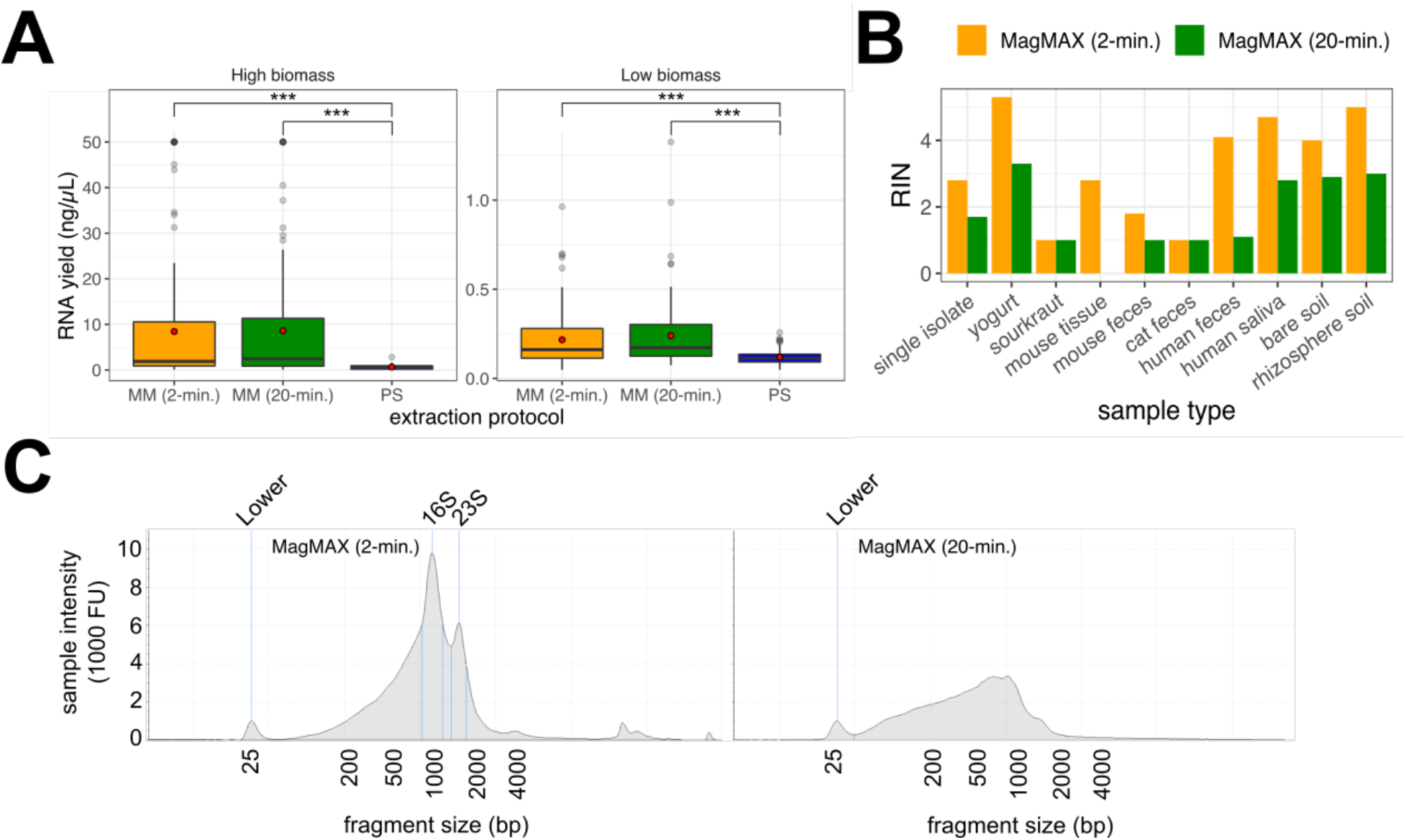
RNA output across extraction protocols. (A) RNA yield quantified using the Qubit RNA assay. Red circles indicate means. Asterisks indicate significant differences between pairs of extraction protocols as determined by paired-sample Wilcoxon signed-rank tests; *** *p* < 2.2E-16. MM = MagMAX; PS = PowerSoil. Values at 50 ng/μL are at the upper limit of detection for the Qubit assay, and may underestimate actual yields for those samples. (B) RNA Integrity Number (RIN) across a subset of samples for the MagMAX extraction protocols, estimated using the TapeStation high-sensitivity (HS) RNA assay. PowerSoil extracts were excluded from the assay due to poor RNA yield, however we note that this may be to our exclusion of the RNAse step available in that protocol. (C) RNA fragment length distribution estimated using the TapeStation HS RNA assay for one human fecal sample. The distribution for the MagMAX (2-min) is on the left and that for the MagMAX (20-min) on the right. The positive control marker at 25-bp is annotated. Peaks corresponding to expected lengths for 16S and 23S rRNA are annotated for the 2-min. protocol and are missing from output from the 2-min. one.

## Conclusions

We conclude that the MagMAX (2-min) extraction protocol is comparable to our established PowerSoil protocol with respect to characterizing microbial community composition, and therefore should allow for comparisons such as meta-analysis across 16S and metagenomics data produced using both protocols, and downstream methods similar to those used here. In addition to extracting both DNA and RNA, the more rapid processing time (*i.e.*, *ca*. 2 h faster than PowerSoil per 96 samples), use of fewer consumables (*i.e.*, *ca* 70% of plastics), and lower cost (*i.e.*, $5.56 vs. $5.65 per sample), highlight the MagMAX (2-min) protocol as an comparable and efficient alternative to the PowerSoil protocol that also allows for downstream applications using RNA.

## Future perspective

Future optimization of molecular methods for microbial community analyses should focus on increasing representation by all microbes in a sample, and from across diverse samples types including those used here. Achieving these goals will allow for more widely adopted use of the same methods. As no single study can be completely comprehensive, making advances that allow us to better compare across studies, particularly with those in the past, is an important step [38]. Alongside the development of computational methods that bioinformatically reduce experimental variation, continuing to explore new molecular methods for capturing important ecological interactions will support our growing understanding of microbial communities.

## Author contributions

JP Shaffer, C Marotz, P Belda-Ferre, RA Salido, CS Carpenter, LS Zaramela, JJ Minich, G Humphrey, AD Swafford, S Miller-Montgomery, and R Knight designed the study; JP Shaffer, C Marotz, P Belda-Ferre, RA Salido, CS Carpenter, LS Zaramela, M Bryant, and G Humphrey provided samples; JP Shaffer, C Marotz, CS Carpenter, and S Fraraccio performed extractions; RA Salido, M Bryant, K Sanders, and G Humphrey performed quality control and sequencing; JP Shaffer, C Marotz, P Belda-Ferre, C Martino, S Wandro, M Estaki, and RA Salido performed data analyses; JP Shaffer wrote the manuscript, with contributions from all authors.

## Acknowledgements

We thank K Dao, R Moranchel, C Nguyen, C Morris, and R Simpson for providing samples; and J DeReus for assistance with data processing.

## Ethical conduct of research

The human subject work conducted here is approved through UCSD IRB#150275.

## Executive summary

1. We compared our established protocol for DNA extraction against two alternative protocols that also extract RNA.
2. We included a diverse panel of sample types ranging from host-associated to environmental.
3. We also included controls for detecting well-to-well contamination and the limit of detection of microbial cells.
4. We observed sample type-specific differences in DNA extraction efficiency among three extraction protocols.
5. Both new kits were similar with respect to RNA extraction efficiency, but varied in RNA quality
6. Sample type and host identity were stronger drivers of microbial community beta-diversity as compared to the extraction protocol used.
7. We identify a protocol that generates both DNA and RNA, and produces data with high similarity to our established protocol with respect to microbial community alpha-diversity, beta-diversity, and taxonomic composition.
8. The similarity between the optimal protocol and our existing one will allow for meta-analyses across both with negligible technical bias.

