## Supplemental materials for "A comparison of DNA/RNA extraction protocols for high-throughput sequencing of microbial communities"

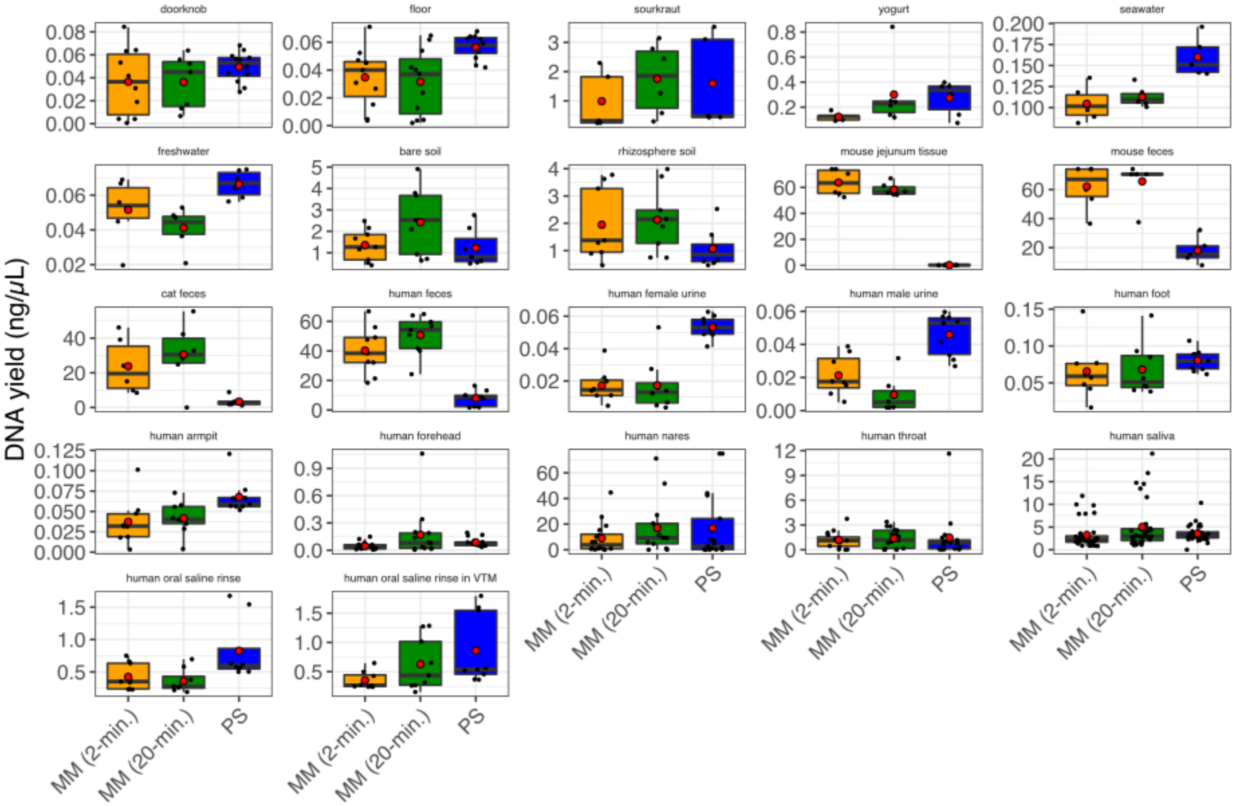


**Figure S1.** Average concentration of DNA (ng/µL) across extraction protocols for each sample type (*n* = 660 total samples included). MM = MagMAX; PS = PowerSoil. Red circles indicate group means. Note that additional samples included here absent from our statistical test (*n* = 45) are those for which technical replication across protocols was not feasible due to recommended sampling protocols (e.g., human nares, human throat), so biological replicates were included instead. Sample types missing here lacked representation by both MagMAX protocols.

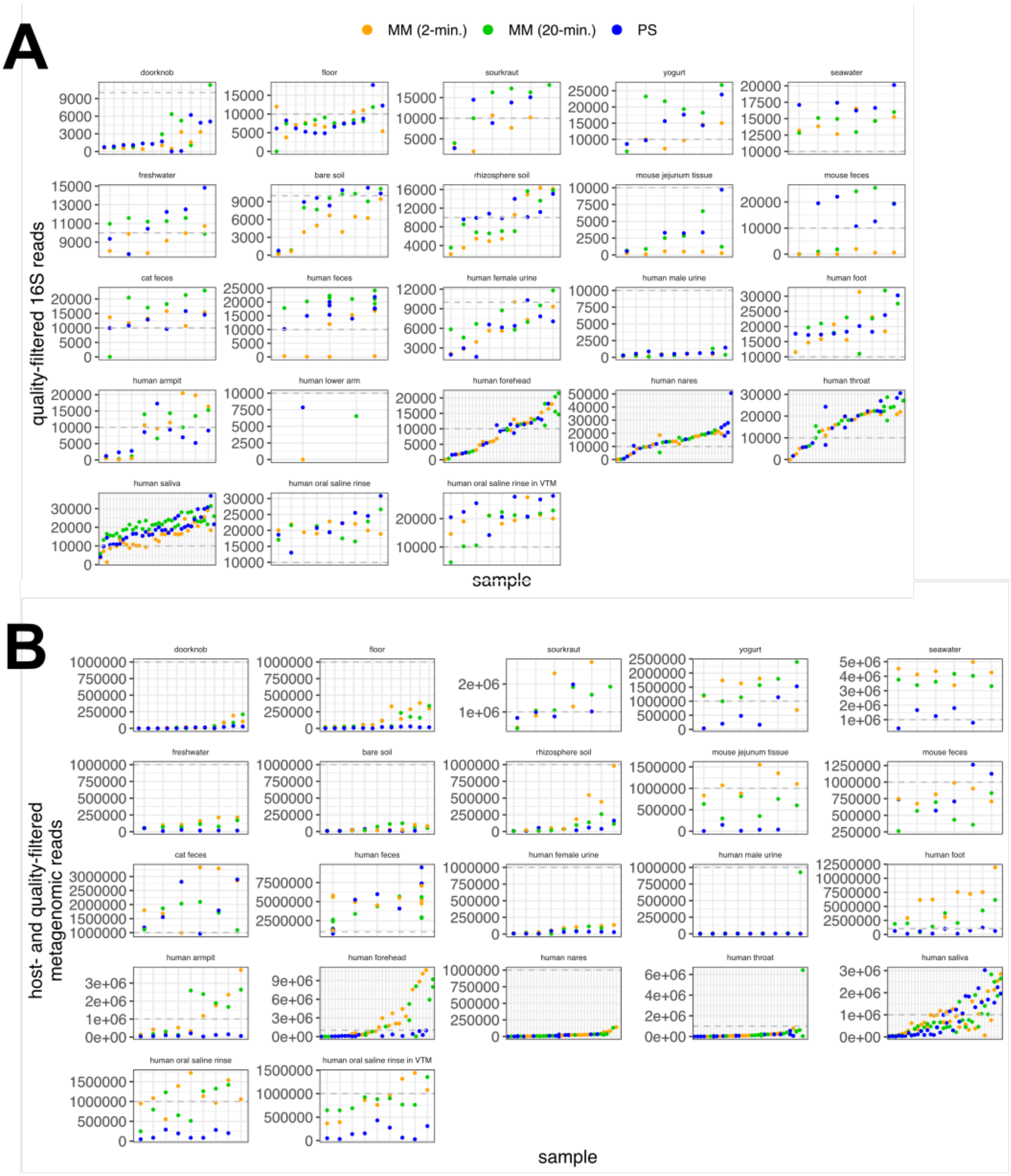


**Figure S2.** Read counts per sample highlighting differences among extraction protocols for each sample type (*n* = 660 samples). (A) Quality-filtered 16S reads for high- and low-biomass samples. (B) Host- and quality-filtered metagenomics reads for high- and low-biomass samples. MM = MagMAX; PS = PowerSoil. Note that additional samples included here absent from our statistical test (*n* = 45) are those for which technical replication across protocols was not feasible due to recommended sampling protocols (e.g., human nares, human throat), so biological replicates were included instead.

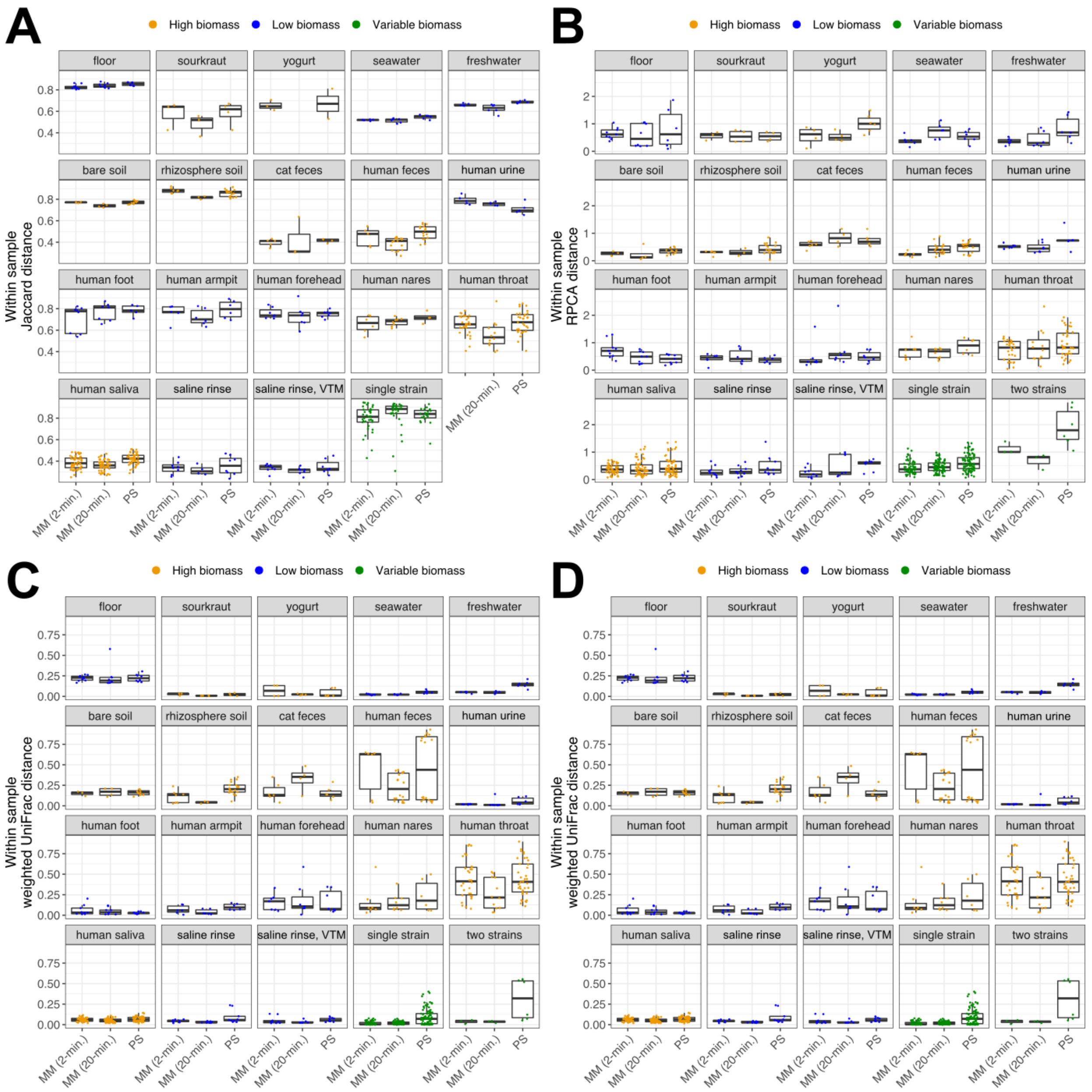


**Figure S3.** Within-sample variation across extraction protocols, for 16S data. Microbial community beta-diversity among replicate extractions of the same source sample was estimated using (A) Jaccard distance, (B) RPCA distance, (C) unweighted UniFrac distance, and (D) weighted UniFrac distance. MM = MagMAX; PS = PowerSoil. Data were rarefied as noted for Figure 3. Sample types missing here lacked representation by both MagMAX protocols.

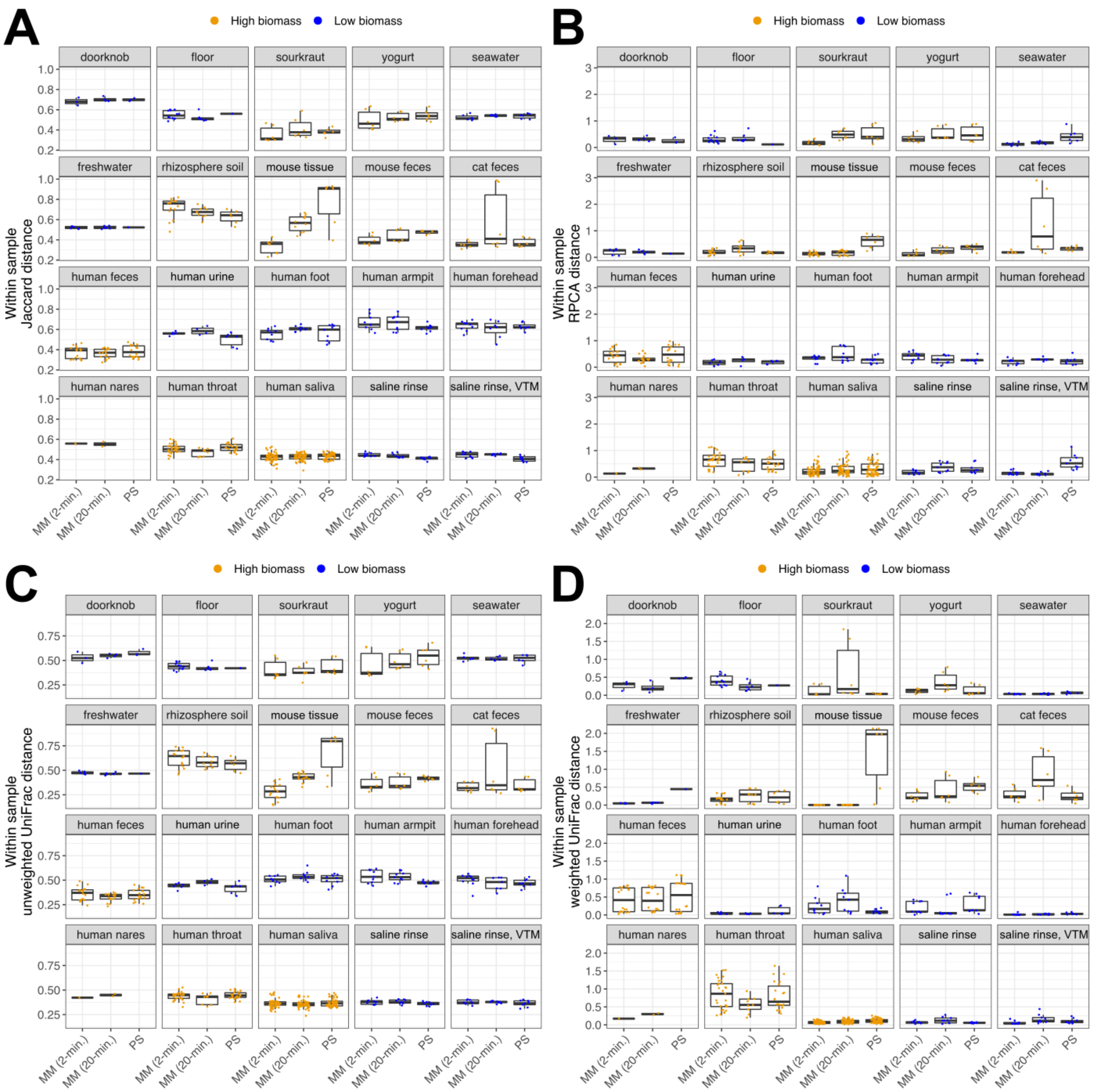


**Figure S4.** Within-sample variation across extraction protocols, for shotgun metagenomics data. Microbial community beta-diversity among replicate extractions of the same source sample was estimated using (A) Jaccard distance, (B) RPCA distance, (C) unweighted UniFrac distance, and (D) weighted UniFrac distance. MM = MagMAX; PS = PowerSoil. Data were rarefied as noted for Figure 3. Sample types missing here lacked representation by both MagMAX protocols.

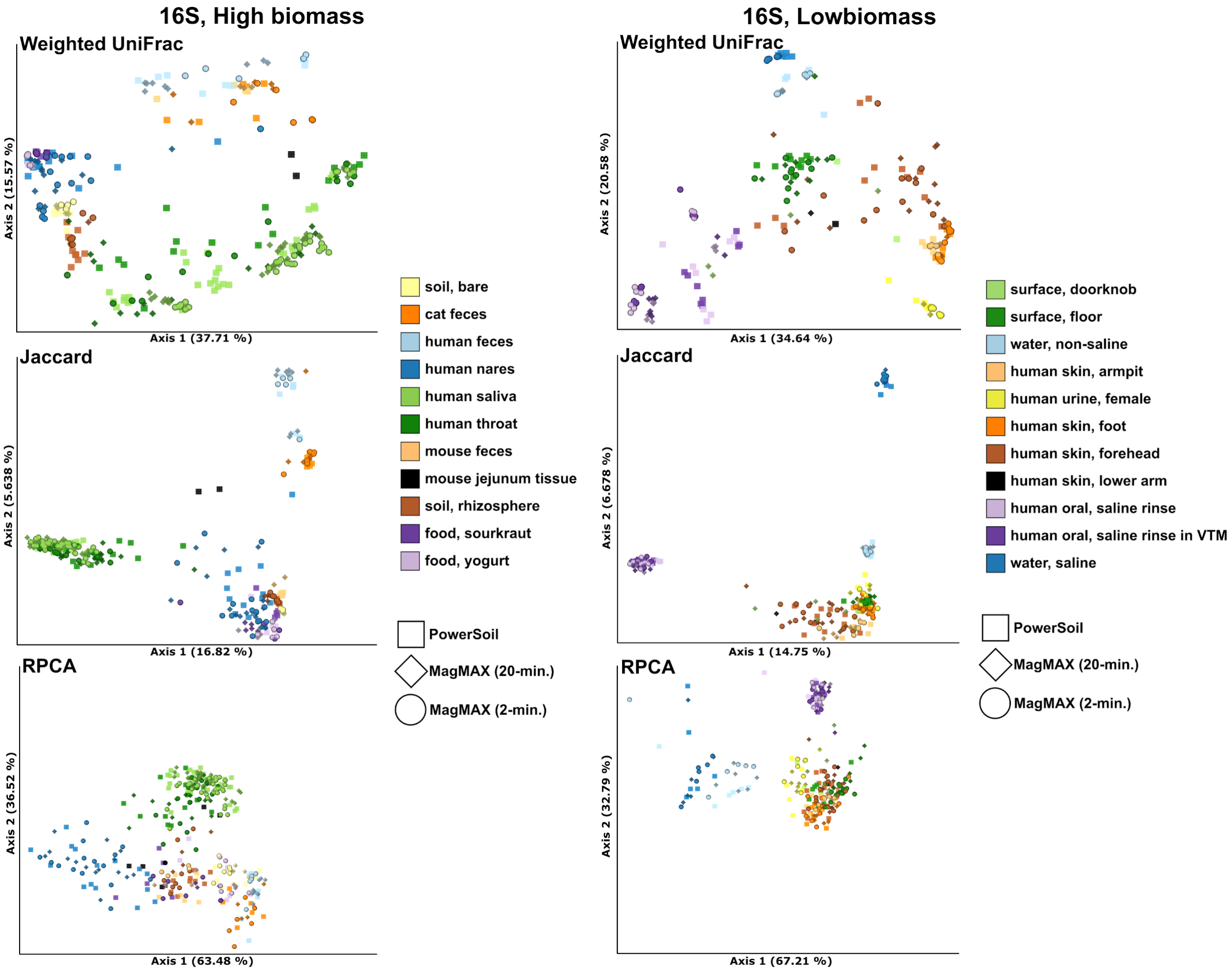


**Figure S5.** Principal coordinates analysis (PCoA) plots showing weighted UniFrac and Jaccard distances, and Principal components analysis (PCA) plots showing RPCA distances, based on 16S data for high- and low-biomass samples. Colors indicate sample types and shapes extraction protocols. Mock community and control blanks were excluded for clarity. Data were rarefied as noted for Figure 3.

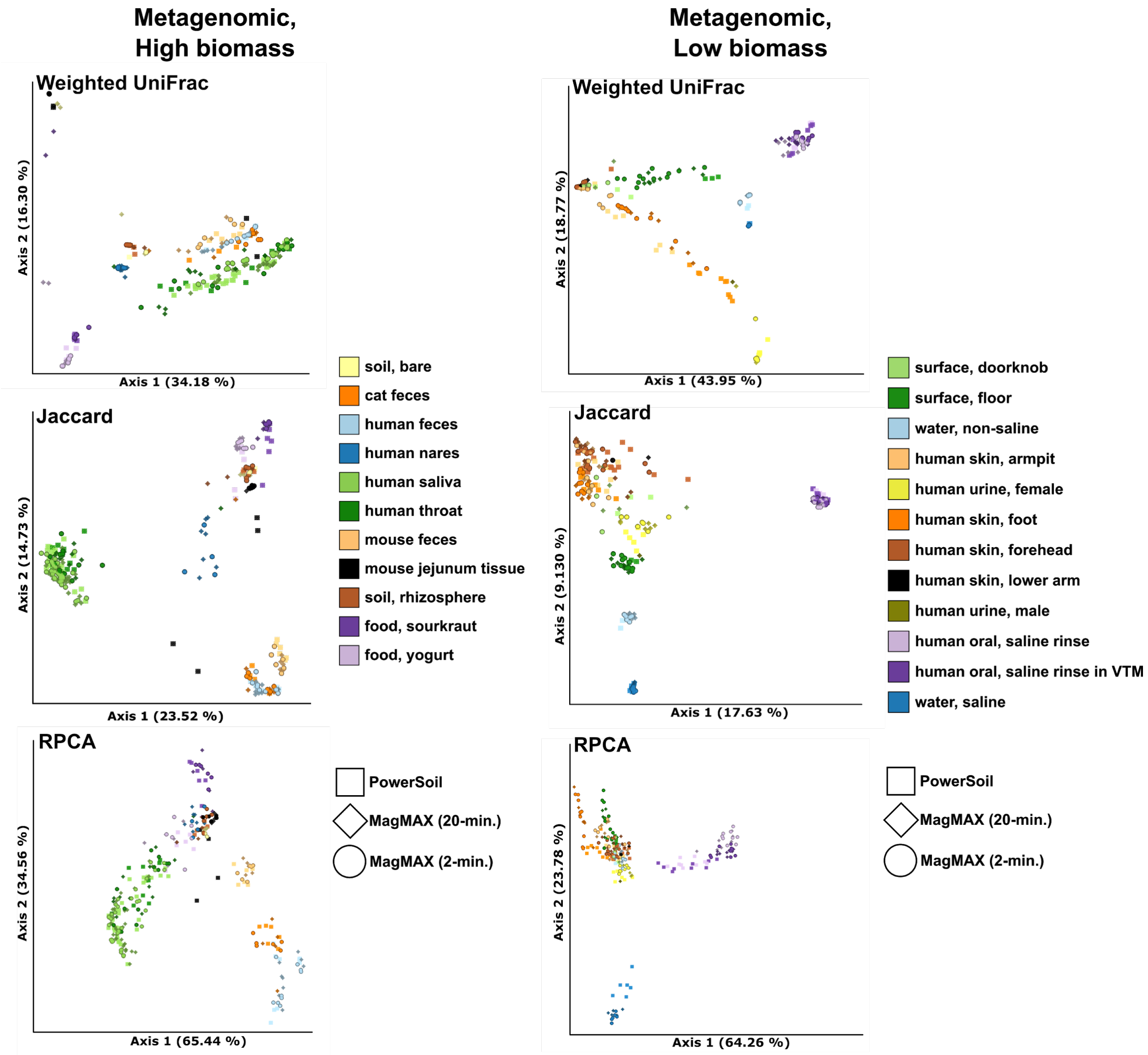


**Figure S6.** Principal coordinates analysis (PCoA) plots showing weighted UniFrac and Jaccard distances, and Principal components analysis (PCA) plots showing RPCA distances, based on shotgun metagenomics data for high- and low-biomass samples. Colors indicate sample types and shapes extraction protocols. Mock community and control blanks were excluded for clarity. Data were rarefied as noted for Figure 3.

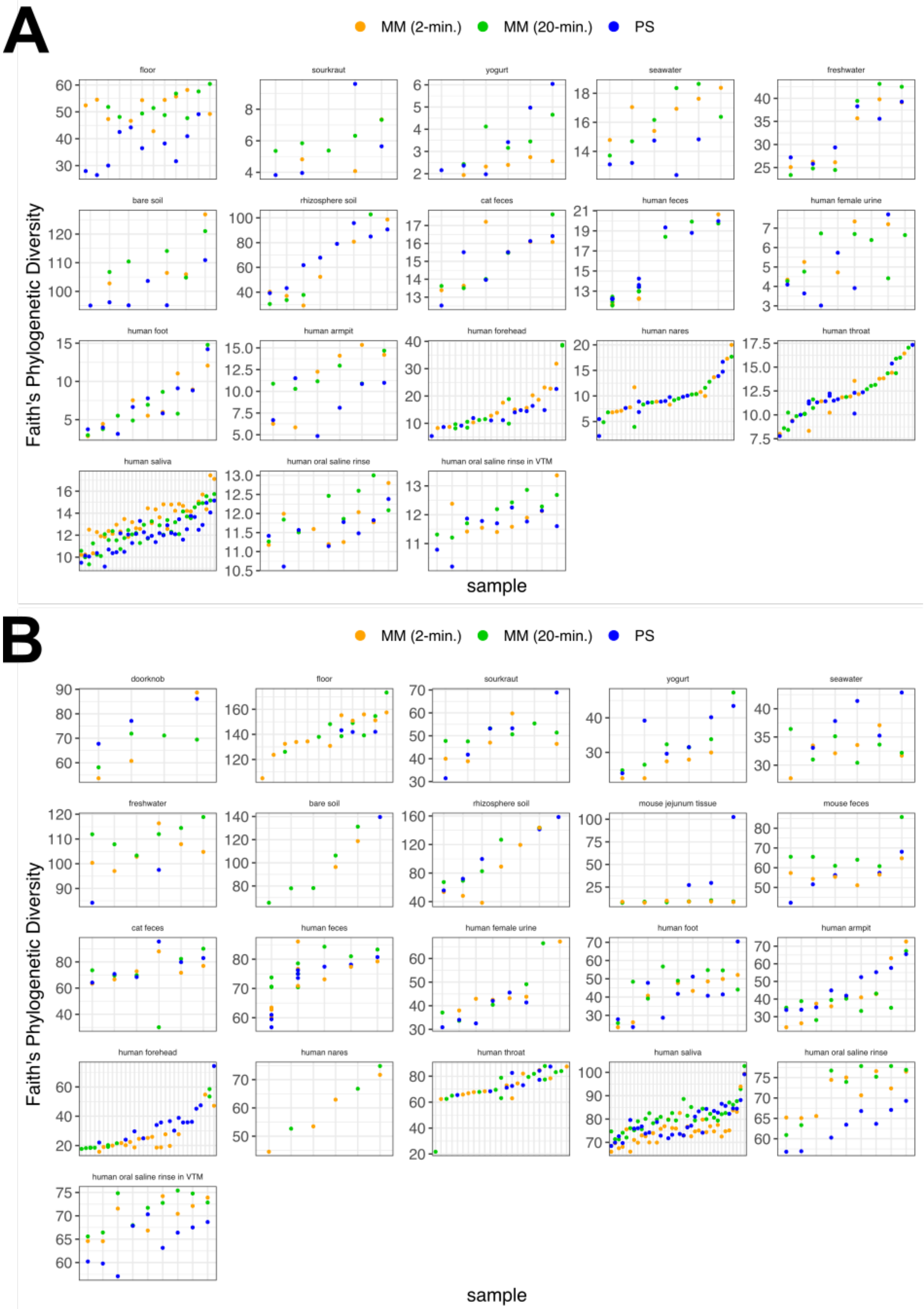


**Figure S7.** Alpha-diversity (Faith's Phylogenetic Diversity) among the three extraction protocols based on (A) 16S and (B) metagenomics data. MM = MagMAX; PS = PowerSoil. Data were rarefied as noted for Figure 3. Sample types missing here lacked representation in at least one of the three extraction protocols.

**Table S1.** Mantel correlations in pairwise distances among samples between all pairs of extraction protocols, for both 16S and metagenomics data. 16S data were rarefied to 5,000 quality-filtered reads per sample, or had samples with fewer than 5,000 reads excluded when using RPCA distances (*n* = 611 samples). Metagenomics data were rarefied to 17,000 host- and quality-filtered reads per sample, or had samples with fewer than 17,000 reads excluded when using RPCA distances (*n* = 647 samples). Rarefaction depths were selected to maintain at least 75% samples from both high- and low-biomass datasets.

| data type | comparison | distance metric | n | Pearson's *r* | p-value |
|---|---|---|---|---|---|
| 16S | MagMAX 2- vs. 20-min. | Jaccard | 137 | 0.96 | 0.002 |
| | | unweighted UniFrac | | 0.92 | 0.002 |
| | | weighted UniFrac | | 0.97 | 0.002 |
| | | RPCA | | 0.92 | 0.002 |
| | MagMAX 2-min. vs. PowerSoil | Jaccard | 139 | 0.96 | 0.002 |
| | | unweighted UniFrac | | 0.93 | 0.002 |
| | | weighted UniFrac | | 0.93 | 0.002 |
| | | RPCA | | 0.90 | 0.002 |
| | MagMAX 20-min. vs. PowerSoil | Jaccard | 159 | 0.95 | 0.002 |
| | | unweighted UniFrac | | 0.93 | 0.002 |
| | | weighted UniFrac | | 0.91 | 0.002 |
| | | RPCA | | 0.89 | 0.002 |
| metagenomic | MagMAX 2- vs. 20-min. | Jaccard | 173 | 0.98 | 0.001 |
| | | unweighted UniFrac | | 0.97 | 0.001 |
| | | weighted UniFrac | | 0.91 | 0.001 |
| | | RPCA | 228 | 0.88 | 0.001 |
| | MagMAX 2-min. vs. PowerSoil | Jaccard | 158 | 0.96 | 0.001 |
| | | unweighted UniFrac | | 0.94 | 0.001 |
| | | weighted UniFrac | | 0.87 | 0.001 |
| | | RPCA | 221 | 0.85 | 0.001 |
| | MagMAX 20-min. vs. PowerSoil | Jaccard | 156 | 0.96 | 0.001 |
| | | unweighted UniFrac | | 0.94 | 0.001 |
| | | weighted UniFrac | | 0.84 | 0.001 |
| | | RPCA | 224 | 0.80 | 0.001 |
